# Dynamics of Innate Immune Response Due to Bacteria-Induced Pulpitis

**DOI:** 10.1101/2023.01.15.524125

**Authors:** Ozge Erdogan, Jingya Xia, Isaac M. Chiu, Jennifer L. Gibbs

**Author notes:** **Corresponding Authors** Jennifer L. Gibbs, Harvard School of Dental Medicine, Department of Restorative Dentistry and Biomaterials Sciences, 188 Longwood Avenue, Boston, MA 02115, Isaac M. Chiu, Harvard Medical School, 77 Avenue Louis Pasteur, Boston, MA 02115. These authors contributed equally (Isaac M. Chiu, Jennifer L. Gibbs).

## Abstract

**Introduction:** Pulpitis is associated with dental carries and can lead to irreversible pulp damage. As bacteria penetrate deeper into dentin and pulp tissue, a pulpal innate immune response is initiated. However, an understanding of the types of immune cells in the pulp, how this relates to bacterial infiltration, and the dynamics of the immune response during pulpitis is limited. As conserving the vitality of the pulp tissue through conservative therapies becomes an important part of dental practice, there is a greater need to understand the kinetics and composition of the immune response during pulpitis.

**Methods:** Dental pulp exposure in molars of mice was used as an animal model of pulpitis. To investigate the kinetics of immune response, pulp tissue was collected from permanent molars at different time points after injury (baseline, day 1, and day 7). Flow cytometry analysis of CD45+ leukoctyes including macrophages, T cells, neutrophils and monocytes was performed. 16S in situ hybridization captured bacterial invasion of the pulp, and immunohistochemistry for F4/80 investigated spatial and morphological changes of macrophages during pulpitis. Data were analyzed using two-way ANOVA with Tukey’s multiple comparisons.

**Results:** Bacteria mostly remained close to the injury site, with some expansion towards non-injured pulp horns. We found that F4/80^+^ macrophages were the main immune cell population in healthy pulp. Upon injury, CD11b^+^Ly6G^high^ neutrophils and CD11b^+^Ly6G^int^Ly6C^int^ monocytes constituted 70-90% of all immune populations up to 7 days after injury. Even though there was a slight increase in T cells at day 7, myeloid cells remained the main drivers of the immune response.

**Conclusions:** As bacteria proliferate within the pulp chamber, innate immune cells including macrophages, neutrophils and monocytes predominate as the major immune populations, with minimal signs of transitioning to an adaptive immune response.

## INTRODUCTION

Non-surgical root canal therapy is a successful and highly predictable treatment option for treating dental pulp inflammation and infection (Burns et al. 2022). However, there has been evidence showing that more conservative treatment options, such as vital pulp therapies, can serve as successful treatment choices and allow reversal of tissue damage (Asgary et al. 2014; European Society of Endodontology developed et al. 2019; Taha and Khazali 2017). There are various challenges and gaps in knowledge in order for clinicians to provide vital pulp therapies as a predictable and reliable treatment option. One key area where there is limited understanding is the dynamics of the immune response and how this correlates with tissue damage and bacterial progression within the coronal pulp tissue.

In the pulp, resident macrophages have been shown to be the main resident immune cell-type that provides the surveillance of the healthy pulp tissue (Gaudin et al. 2015; Krivanek et al. 2020). However, in the context of infection and pulpitis, less is known about the kinetics of the immune response, mainly because the analysis of these changes are technically challenging (Galler et al. 2021). Myeloid immune cells, which include macrophages, neutrophils and monocytes are crucial members of the innate immunity. Neutrophils and monocytes are circulating leukoctyes that play a critical role in antibacterial host defense, and are one of the first innate immune cells recruited to the site of infection or injury (de Oliveira et al. 2016; Perez-Figueroa et al. 2021; Segal 2005). Neutrophils ingest and phagocytose bacteria, and their entry into tissues and subsequent cell death sets the stage for the resolution phase of the inflammation or for a successful transition to adaptive immune response (Kolaczkowska and Kubes 2013; Rosales 2020). A dysregulation of neutrophil recruitment and function can contribute to continuous tissue damage as seen in some of chronic inflammatory diseases (Chamardani and Amiritavassoli 2022; Schultz et al. 2022). Previous studies have also investigated a possible destructive role of neutrophils in the pulp tissue (Holder et al. 2019; Nakamura et al. 2002). Understanding the dynamics of these immune cell populations during pulpitis can provide insight about the protective and destructive processes occurring during pulpitis (Duncan and Cooper 2020; Renard et al. 2016).

Past studies examining human pulp tissue samples have investigated how the innate and adaptive immune response may relate to microbiome changes in dental caries (Sakurai et al. 1999; Yoshiba et al. 2003; Yoshiba et al. 2020; Yoshiba et al. 1996). However, these studies can only capture one time point, rather than the dynamics of the response, and are limited to human samples collected once disease has progressed significantly that the teeth require extraction or root canal treatment. Therefore, studying the innate immune response which occurs at the initial stages of pulpal inflammation is challenging in human tissues.

Using animal models of pulpitis would enables a better capture of the dynamics of the cellular response. A previous study using an injury model of mouse incisors showed that there was a major increase in the percentage of myeloid cells after injury which almost returned to baseline by three days after injury (Renard et al. 2016). However, in this study, the investigators used murine incisor teeth, which grow continuously even after the teeth complete their maturation (Krivanek et al. 2020). Therefore, it is possible that the immune response of continuously growing mice incisor teeth would differ than human teeth. In this study, we aimed to use a mouse model of pulpitis based on molar tooth injury which may more accurately reflect human pulpitis.

We applied flow cytometry of the pulp to perform a detailed investigation of the kinetics of the immune cell response during pulpitis in mice. We correlated this with histological analysis of bacterial infection. Spatially and kinetically we found that macrophages were the resident immune cells and they showed morphological changes that corresponded with the timeline of bacterial invasion. We also found that neutrophils and monocytes were the main innate immune cell populations that entered the pulp and persisted until later stages of the disease.

## MATERIALS AND METHODS

### Animals

All animal experiments were approved by the Institutional Animal Care and Use Committee (IACUC) at Harvard Medical School. C57BL/6 mice were purchased from Jackson (Jax) Laboratories (Bar Harbor, ME). Age-matched 7-to 12-week-old littermate mice of both genders were used for experiments.

### Dental Pulp Injury Procedure

Mice were anesthetized via intraperitoneal injection of 100 mg/kg ketamine + 10 mg/kg xylazine in sterile phosphate buffered saline (PBS). The exposures were generated in accordance with published protocols (Gibbs et al. 2013; Lee et al. 2017). Briefly, the molar tooth was drilled with a ¼ round bur at low speed until the pulp was exposed. The contralateral molar served as the uninjured control. For histology experiments, only the maxillary first molar for each animal was injured. For flow cytometry experiments, maxillary first, second and mandibular first molars were injured in each animal and teeth from 5 mice were pooled for one sample.

### Pulp Tissue Cell Suspension Preparation

Maxillae/and mandibles were dissected, put in minimum essential medium (MEM-α) on ice. Surrounding tissues of teeth, including gingiva were removed. The crowns of teeth were separated using a scalpel blade at crown-root interference which allowed both for exposure and collection of coronal pulp tissue by avoiding any contamination from surrounding tissues. The crowns were placed into the MEM-α culture media. Culture media was then discared and the crowns were incubated in digestive solution (4mg/ml collagenase and 4mg/ml dispase II) at 37 °C for 1 hour. Later, the crowns within the digestive solution were pipetted vigorously for the pulp tissue to be detached from dentin, the samples were filtered through using 70 μm mesh. The pulp samples were spun down at 300g for 5 minutes, washed with flow buffer (2% FBS, 1μM EDTA, 0.1% sodium azide).

### Flow Cytometry

Cell suspensions were spun down at 300g for 5 minutes and resuspended in flow buffer. The cell suspension was incubated with FcR blocking reagent for ten minutes, washed with flow buffer and then incubated with the following antibodies (Zombi Aqua -KO525 (1:1000, Biolegend), CD45-Cy7 (1:100, Biolegend), CD11b-PE (1:100, Biolegend), Ly6G-APC (1:100, Biolegend), Ly6C-AF700 (1:100, Biolegend), CD3-FITC (1:100, Biolegend), CD4-PE-cy7 (1:100, Biolegend), CD8-perpCy5.5 (1:100, Biolegend), F4/80-PB (1:100, Biolegend) for 30 minutes. The cell suspensions were washed again, centrifuged, resuspended in 1% paraformaldehyde (PFA) and filtered through a 40 μm mesh. Counting beads (Biolegend) were added to each sample. Flow cytometry was performed on a CytoFLEX flow cytometer (Beckman Coulter Life Sciences) and data was analyzed using FlowJo software (FlowJo LLC).

### Tissue Processing & Immunofluorescence Staining

Animals were perfused with PBS with heparin, followed by 4% PFA. Maxillae were dissected out and cut out to separate right (injured tooth side) and left (non-injured tooth side) half maxillae. All tissues were post-fixed with 4% PFA. Decalcification of maxillae using 18% EDTA for 3-4 weeks at 4°C was performed. Half maxillae were dehydrated by 30% sucrose for 2 days, followed by 50% sucrose for 1 day at 4°C and then cryopreserved. 30µm cryosections were obtained. Cryosections were washed with PBS for 30 minutes. Sections were blocked with 5% goat serum and 1% triton for 1 hour at room temperature. Sections then were incubated with the primary antibody -rat anti-F4/80 (1:200, abcam)-overnight at 4°C. Sections were then washed with 1% goat serum and 1% triton for 30 minutes and were incubated with the secondary antibody -anti-rat Alexa fluor647 (1:500, ThermoFisher)-for 2 hours. Stained slides were mounted. Image acquisition was performed by confocal microscope (Leica, Stellaris). Equal number of stacks were imaged using the same acquisition settings. Image processing was performed using Fiji (Schindelin et al. 2012) by using the same settings.

### Fluorescence In-situ Hybridization

Cryosections were washed with PBS for 30 minutes. Half of the sections on one slide were incubated with 1:200 Cy3-congugated nonsense eubacterial probe (5’-/5Cy3/CGACGGAGGGCATCCTCA-3’) whereas the other half of the sections on the same slide were incubated with 1:200 Cy3-conjugated 16S eubacterial probe (5’-/5Cy3/GCTGCCTCCCGTAGGAGT-3’) in 20% SDS, 1:10 formamide, and 9:10 bacterial hybridization buffer overnight in 50°C in hybridization oven. Sections were washed first with hybridization buffer followed by PBS before mounting.

### Statistical Analysis

Statistical comparisons of two groups dependent on a single variable with normal distributions were analyzed by an unpaired t test. Statistical comparisons of three or more groups at a single time point was analyzed by one-way ANOVA with Tukey post-tests. Between subject effects were used to determine if there had been any significant effects of time, experimental group, or a significant interaction, followed by with post-hoc Dunnett’s and Sidak tests where necessary. A p-value less than 0.05 was considered statistically significant. Statistical analyses were conducted using Prism 9 (GraphPad Software, LLC) and Excel (Microsoft).

## RESULTS

### Bacteria remain within the coronal pulp tissue up to day seven during pulpitis

We wanted to first observe the penetration of bacteria into the pulp space after pulp exposure over the same time frame that we captured the immune response. By performing florescence in situ hybridization (FISH) of 16S bacterial rRNA in tissue sections, we found that at day 0, just after the injury, there were no bacteria present at the injury site (Figure 1-A). At day 1 post-injury, bacteria were clearly present at the injury site, superficially located at the injured middle pulp horn (Figure 1-B). By day 3 and 7, the majority of bacteria were still densely present close to the injury site. However, we also observed bacteria spreading to non-injured pulp horns and even at the coronal third of the root pulp (Figure 1-C, D).

**Figure 1.**
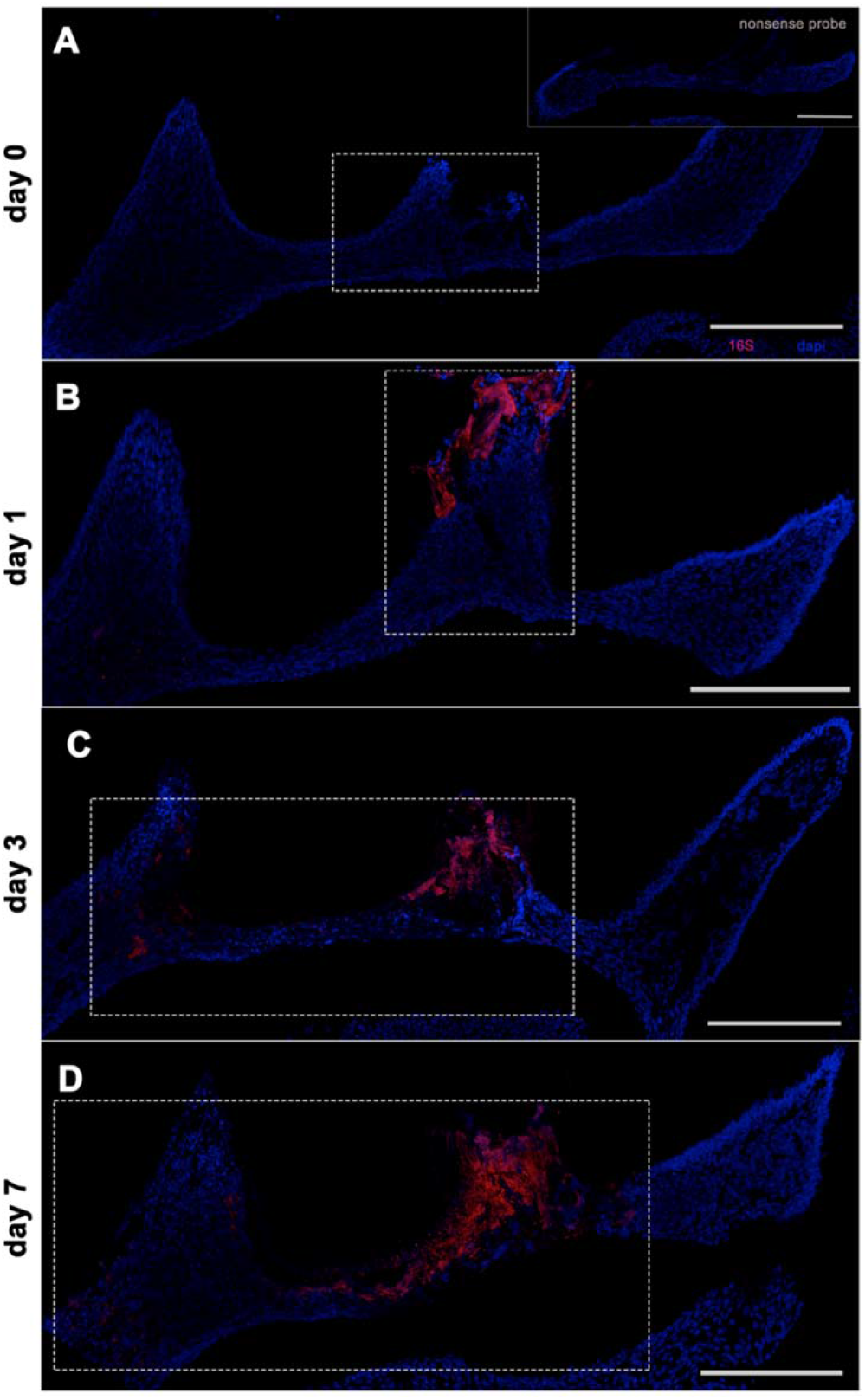
16S florescence-in situ hybridization to investigate bacteria localization within the pulp chamber across different time points. A- Pulp, just right after injury. The dashed box shows the mechanically injured middle pulp with no bacteria present. The image on the top right corner is a section where the nonsense probe was used as a negative control. B- Pulp, one day after injury. The bacteria are localized at the injury site, shown within the box. C, D- Pulp, three and seven days after injury. The boxes show that most of the bacteria are still present at the injury site but extended to the non-injured pulp horn. Scale bars are 200 μm. Red-16S probe, blue-dapi.

### Immune cell populations of healthy pulp tissue

We first wanted to study immune survelience of healthy pulp tissue. Histologically, we observed that F4/80+ macrophage population was densely present in the healthy pulp tissue (Figure 2-A). They exhibited an amoeboid cell shape with numerous cytoplasmic extensions (Figure 2-A). We next performed flow cytometric analysis of different subsets of CD45+ immune cells in the healthy dental pulp including macrophages, neutrophils, monocytes, and T cells. In healthy intact pulp tissue, CD45+ immune cells constituted on average of only 3% of all viable cells (Figure 3-B, F). CD11b+ myeloid cells were the major immune cell population (74%) present, whereas only about 2.6% of resident immune cells were CD3+ T cells (Figure 3-C, G, H). Of CD11b+ myeloid cells, F4/80+ macrophages were the main immune cell population, forming on average 68% of all immune cells (Figure 3-D H).

**Figure 2.**
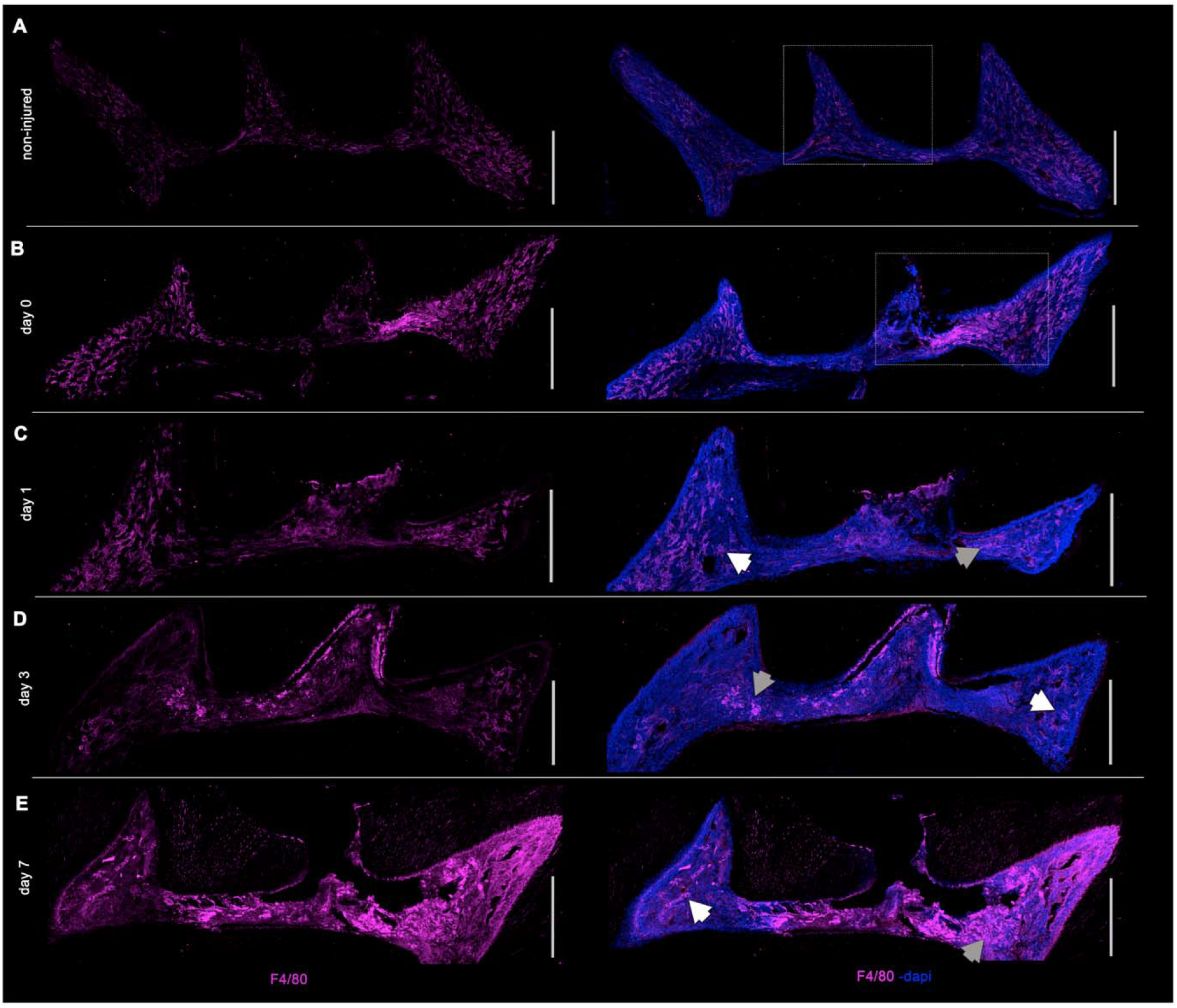
Spatial and morphological changes of F4/80+ macrophages across different time points within the pulp chamber. A- Non-injured pulp. The box shows the intact middle pulp horn with densely and evenly located macrophages. B- Pulp, just right after injury. The box shows the injured middle pulp horn where macrophages are not visible and a part of non-injured pulp horn where macrophages with their distinct morphology are present. C, D, E- Pulp, one day, three days and seven days after injury. The white arrows point out the morphologically intact macrophages within the non-injured pulp horn whereas the gray arrows point out an area where the morphology of the macrophages look less distinct. The middle pulp horn is the mechanically injured pulp horn in each image. The first column includes images showing F4/80+ immunofluorescence, the second column shows F4/80 co-localization with dapi. Scale bars are 200 μm.

**Figure 3.**
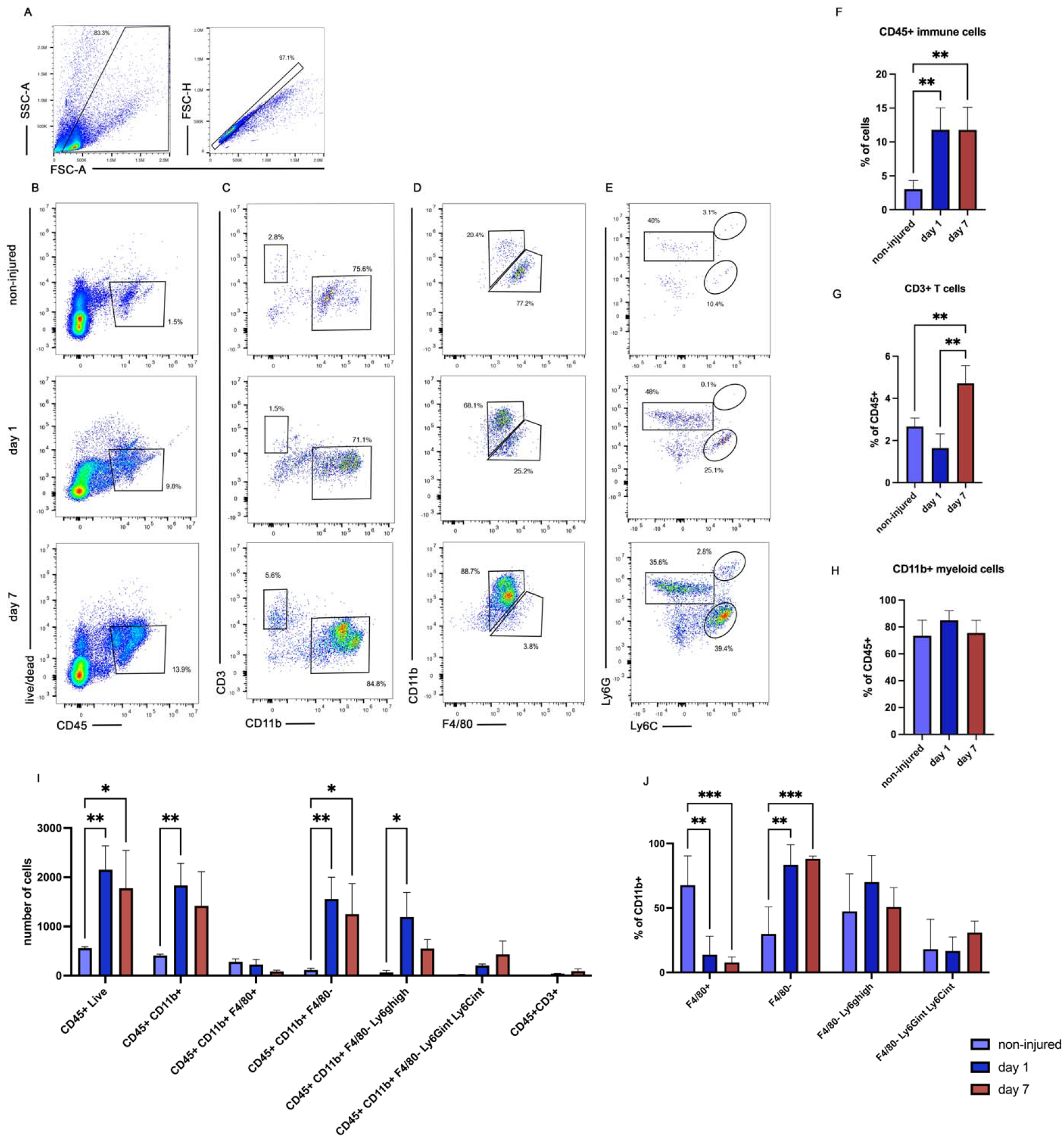
Myeloid cells, especially neutrophils are the major immune cell populations up to 7 days after pulp injury. A- Representative flow cytometry (FC) plots showing initial steps of the gating strategy. B- Representative FC plots to gate CD45+ live immune cells of non-injured pulp, injured pulp one and seven days after injury. C- Representative FC plots to gate CD3+ T cells and CD11b+ myeloid cells of non-injured pulp, injured pulp one and seven days after injury. D- Representative FC plots to gate F4/80+ macrophages of non-injured pulp, injured pulp one and seven days after injury. E- Representative FC plots to gate F4/80- myeloid cells into Ly6Ghigh, Ly6Gint Ly6Cint and Ly6Ghigh Ly6Chigh neutrophils/monocytes of non-injured pulp, injured pulp one and seven days after injury. F- % of CD45+ immune cells among all cells at different time points. G- % of CD3+ T cells among CD45+ cells at different time points. H- % of CD11b+ myeloid cells among CD45+ immune cells at different time points. I- Number of immune cell changes plotted from three independent experiments. J- % of different myeloid cell populations (macrophages, neutrophils and monocytes) at different time points. I- Two-way ANOVA with Tukey’s multiple comparisons test. (type of immune cells: p<0.0001, F=10.25, DOF=6; time: p<0.0001 F=14.79, DOF=2; interaction: p=0.07, F=1.7, DOF=12). J- Two-way ANOVA with Dunnett’s multiple comparisons test (type of immune cells: p<0.0001, F=13, DOF=3; time: p=0.7 F=0.3, DOF=2; interaction: p<0.0001, F=7.9, DOF=6). F, G, H- One-way ANOVA with Tukey’s multiple comparisons test ∗p < 0.05, ∗∗p < 0.01, ∗∗∗∗p < 0.0001. Mean ± SEM.

### F4/80+ macrophages are resident and respond to pulp injury

Histologically and kinetically, we studied macrophages in response to injury. On day 1, 3 and 7 after injury, the morphology of macrophages in non-injured areas of the pulp was similar to healthy pulp tissue with amoeboid cell shape with numerous cytoplasmic extensions present (Figure 2-B, C, D). However, histologically, on day 3 after injury in the areas close to the injury site and on day 7 within the whole coronal pulp, macrophages exhibited an enlarged, amoeboid shape with fewer cytoplasmic extensions (Figure 2-D, E). The timing and location of these morphological changes of macrophages corresponded well with the expansion timeline of the bacteria. We also studied the changes in F4/80+ macrophages quantitatively in response to injury, given that they were the main resident immune cell population at baseline. Out of total CD45+ immune cells, the total percentages of F4/80+ macrophages decreased at day 1 (p=0.001) and 7 (p=0.001) compared to healthy pulp, which is likely due to the concomitant increased proportions of neutrophils and monocytes (Fig. 3-J). However, by cell count, the total number of F4/80+ cells remained the same across all time points in pulp tissues (Figure 3-I). These findings indicate that macrophages are the main resident immune cells of the pulp tissue and respond to injury.

### Cd11b+ myeloid cells, especially neutrophils are the major immune cell populations up to seven days after pulp injury

Upon injury, compared to healthy pulp tissue, the percentage and total number of CD45+ immune cells increased and constituted about 12% of all viable pulp tissue cells at both day 1 and day 7 (Figure 3-B, F). The total number of CD45+ leukocytes also increased significantly at day 1 (p=0.001) and at day 7 compared to baseline (p=0.015) (Figure 3-I). Significant changes occurred within the myeloid immune cell population. Even though the percentage of CD11b+ myeloid cells out of total immune cells remained similar at day 1 and 7, their numbers increased significantly compared to healthy pulp (comparision of baseline to day 1; p=0.003,baseline to day 7; p=0.051) (Figure 3-H, I). The composition of this myeloid cell population also changed. In the healthy pulp tissue, most of the myeloid cells were F4/80+ macrophages. Upon injury, the major myeloid cell populations became CD11b+Ly6G^high^ neutrophils and CD11b+Ly6G^int^Ly6C^int^ monocytes (Figure 3-D, E, J). Among these neutrophil and monocyte populations, both the percentage and numbers increased significantly compared to healthy pulp and were similar at day 1 and day 7 after injury (comparision of percentages baseline to day 1; p=0.0001, baseline to day 7; p=0001, comparision of numbers baseline to day ; p=0.003, baseline to day 7; p=0.025) (Figure 3-E, I, J). We did observe that CD3+ T cells significantly increased by day 7 post-injury from 2% at baseline to 4% of total immune cells (Figure 3-G).

## DISCUSSION

To our knowledge, we have described for the first time the dynamic changes in immune responses up to day seven in mice in the molar dental pulp injury model using both flow cytometry and histological analysis. We found that macrophages were the most abundant immune cell population in healthy molars, which is in line with a recent transcriptomics analysis of healthy mice and human molar teeth (Krivanek et al. 2020). We also showed the spatial distribution of these macrophages by performing immunostaining. Their morphology and distribution in healthy pulp tissue resemble previous reports of antigen presenting cells including dendritic cells identified by antibodies in human tissue or mouse lines (Hong et al. 2019; Sakurai et al. 1999; Yoshiba et al. 2003). It is possible that, these resident immune cells are F4/80+, which is the marker we used to identify resident macrophages, and cd11c+, which was the marker that had been used to identify dendritic cell populations (Cao et al. 2015). Another study also described CX3CR1+ macrophages with similar morphology histologically in the pulp tissue (Hong et al. 2019). There is still limited understanding about how these resident macrophages behave during pulpitis and their primary role in orchestrating the immune response. Following injury, we found that F4/80+ macrophages adopted an amoeboid cell shape where cytoplasmic extensions were not as clearly visible. This change in morphology could be due to functional changes including phagocytosis of bacteria or debris (Davies et al. 2013). A recent study investigated the role of macrophages in dentin formation in a pulp capping model, finding that when macrophages were depleted by clodronate liposome injections, there was reduced dentin formation (Neves et al. 2020). Further functional studies are needed to understand macrophage behavior during pulpitis to understand how they drive innate immune response, including whether they play a role in bacterial clearance, and recruitment of subsequent innate or adaptive immune cells.

Upon injury, we found that neutrophils and monocytes were the main immune cell populations recruited on day one. Neutrophils in particular were strongly recruited and remained as a prominent immune cell population up to day seven after injury. The kinetics of this neutrophil recruitment is interesting. In another study using a ligature-induced periodontitis model, it was shown that neutrophils expand on day one following inflammation, but both the percentage and the number of neutrophils decreased after that time point (Alvarez et al. 2021). It is possible that our findings of the continued presence of neutrophils in the pulpitis model may relate to the need to control bacterial invasion. In a skin incision-wound healing model, it was shown that after day two, there was a gradual decrease in number of neutrophils at the wound site (Kim et al. 2008). By contrast, when the incision site was inoculated with the bacterial pathogen *S. aureus*, neutrophils increased twofold compared to vehicle group up to six days after bacterial inoculation (Kim et al. 2008). This suggests that a consistent bacterial presence and release of pathogen-associated-molecular-patterns (PAMPS) may induce a longer lasting neutrophil recruitment to the infected site (de Oliveira et al. 2016). In our pulpitis model, we observed that bacteria presence was expanded where they penetrated deeper into the pulp tissue over time, and this could lead to a longer neutrophil presence.

Our flow cytometry analysis showed that on day seven after injury, there was continuity of acute inflammation and predominately innate immune responses with little evidence for a transition to adaptive immune responses and chronic inflammation. There is also discussion on whether human dental pulp tissue has its lymphatic system and how it functions (Bernick 1977; Gerli et al. 2010; Wisniewska et al. 2021), raising questions about how the adaptive immune response might be also different in the pulp tissue compared other tissues. We did find that there was an increase in T cell population on day 7. There is also evidence of T and B cell presence in inflamed pulp due to caries in studies investigating pulp tissue samples from human teeth (Hahn et al. 1989). Of note, the model we are using in mice expedites the process of a much slower disease progression in humans during pulpitis due to caries. Also, there could be species specific differences in terms of the role of innate and adaptive immune cells in pulpitis. Future studies will be needed to generally dissect the role of distinct innate and adaptive immune populations in pulpitis pathogenesis.

## CONCLUSION

Macrophages are the main immune cell population in healthy pulp tissue, and their cellular morphology changes during pulpitis. By flow cytometry analysis, we found that there is rapid recruitment of neutrophils and monocytes during the early stages following injury, with persistent presence of neutrophils throughout the time course of analysis up to day 7. Histological analysis showed that while most of the bacteria was primarily localized at the injury site, by the end point they begin to spreading out within the pulp chamber. There is minimal transitioning into adaptive immune responses after bacteria-induced pulpitis in mice molars. Therefore, a dynamic innate immune response accompanies bacterial invasion of the pulp in a mouse model of pulpitis.

